# BioDynaMo: a general platform for scalable agent-based simulation

**DOI:** 10.1101/2020.06.08.139949

**Authors:** Lukas Breitwieser, Ahmad Hesam, Jean de Montigny, Vasileios Vavourakis, Alexandros Iosif, Jack Jennings, Marcus Kaiser, Marco Manca, Alberto Di Meglio, Zaid Al-Ars, Fons Rademakers, Onur Mutlu, Roman Bauer

**Affiliations:** CERN openlab, CERN, European Organization for Nuclear Research, Geneva, Switzerland; ETH Zurich, Swiss Federal Institute of Technology in Zurich, Zurich, Switzerland; Delft University of Technology, Delft, the Netherlands; Department of Mechanical & Manufacturing Engineering, University of Cyprus, Nicosia, Cyprus; Department of Medical Physics & Biomedical Engineering, University College London, London, UK; School of Computing, Newcastle University, Newcastle upon Tyne, UK; Department of Functional Neurosurgery, Ruijin Hospital, Shanghai Jiao Tong University School of Medicine, Shanghai, China; Precision Imaging Beacon, School of Medicine, University of Nottingham, Nottingham, NG7 2UH; SCimPulse Foundation, Geleen, Netherlands; Department of Computer Science, University of Surrey, Guildford, UK

## Abstract

**Motivation:** Agent-based modeling is an indispensable tool for studying complex biological systems. However, existing simulators do not always take full advantage of modern hardware and often have a field-specific software design.

**Results:** We present a novel simulation platform called BioDynaMo that alleviates both of these problems. BioDynaMo features a general-purpose and high-performance simulation engine. We demonstrate that BioDynaMo can be used to simulate use cases in: neuroscience, oncology, and epidemiology. For each use case we validate our findings with experimental data or an analytical solution. Our performance results show that BioDynaMo performs up to three orders of magnitude faster than the state-of-the-art baseline. This improvement makes it feasible to simulate each use case with one billion agents on a single server, showcasing the potential BioDynaMo has for computational biology research.

**Availability:** BioDynaMo is an open-source project under the Apache 2.0 license and is available at www.biodynamo.org. Instructions to reproduce the results are available in supplementary information.

**Supplementary information:** Available at https://doi.org/10.5281/zenodo.4501515.

## 1 Introduction

Agent-based simulation is a powerful tool assisting life scientists in better understanding complex biological systems. In silico simulation is an inexpensive and efficient way to rapidly test hypotheses about the (patho)physiology of cellular populations, tissues, organs, or entire organisms (Yankeelov *et al.*, 2016; Ji *et al.*, 2017).

However, the effectiveness of such computer simulations for scientific research is often limited, mainly because of two reasons. First, after the slowing down of Moore’s law (Moore, 1965) and Dennard scaling (Dennard *et al.*, 1974), hardware has become increasingly parallel and heterogeneous. Most simulators do not take full advantage of these hardware enhancements. The resulting limited computational power forces life scientists to compromise either on the resolution of the model or on simulation size (Thorne *et al.*, 2007). Second, existing simulators have often been developed with a specific use case in mind. This makes it challenging to implement the desired model, even if it deviates only slightly from its original purpose.

To help researchers tackle these two major challenges, we propose a novel open-source platform for biology dynamics modeling, BioDynaMo. We alleviate both of these problems by emphasizing performance and modularity. BioDynaMo features a high-performance simulation engine which is fully parallelized and able to offload computation to hardware accelerators. The software comprises a set of fundamental biological functions, and a flexible design that adapts to specific user requirements. Currently, BioDynaMo implements the biological model presented in Zubler and Douglas (2009), but this model can easily be extended, modified, or replaced. Hence, BioDynaMo is well-suited for simulating a wide range of biological processes including cell proliferation, migration, growth, etc.

BioDynaMo provides by design five system properties:

- **Agent-based.** The BioDynaMo project is established to support developmental simulations of biological dynamics. A good approximation for such in silico simulations is agent-based modeling (Railsback and Grimm, 2019). Agents are modeled as discrete objects that perform actions based on their current state, behavior, and the surrounding environment.
- **General purpose.** BioDynaMo is developed to become a general-purpose tool for agent-based simulation. To simulate models from various fields, BioDynaMo’s software design is extensible and modular.
- **Large scale.** Biological systems contain a large number of agents. The cerebral cortex, for example, comprises approximately 16 billion neurons (Azevedo *et al.*, 2009). Biologists should not be limited by the number of agents within a simulation. Consequently, BioDynaMo is designed to take full advantage of modern hardware and use memory efficiently to scale up simulations.
- **Easily programmable.** The success of a simulator depends, among other things, on how quickly a scientist, not necessarily an expert in computer science or high-performance programming, can translate an idea into a simulation. This characteristic can be broken down into four key requirements that BioDynaMo is designed to fullfil: First, BioDynaMo provides a wide range of common functionalities such as visualization, plotting, parameter parsing, backups, etc. Second, BioDynaMo provides simulation primitives that minimize the programming effort necessary to build a use case. Third, as outlined in item “General purpose”, BioDynaMo has a modular and extensible design. Fourth, BioDynaMo provides a coherent API and hides implementation details that are irrelevant for a computational model (e.g., details such as parallelization strategy, synchronization, load balancing, or hardware optimizations).
- **Quality assured.** BioDynaMo establishes a rigorous, test-driven development process to foster correctness, maintainability of the codebase, and reproducibility of results.

The main contribution of this paper is an open-source, high-performance, and general-purpose simulation platform for agent-based simulations. We provide the following evidence to support this claim: (i) We detail the user-facing features of BioDynaMo that enable users to build a simulation based on predefined building blocks and to define a model tailored to their needs. (ii) We present three basic use cases in the field of neuroscience, oncology, and epidemiology to demonstrate BioDynaMo’s capabilities and modularity. (iii) We show that BioDynaMo can produce biologically-meaningful simulation results by validating these use cases against experimental data, or an analytical solution. (iv) We present performance data on different systems and scale each use case to one billion agents to demonstrate BioDynaMo’s performance.

### 1.1 Prior work

The history of agent-based modeling and simulation goes well before the 1990s; however, it has seen widespread use in biological systems in the 2000s. Several simulators have been published demonstrating the importance of agent-based models in computational biology research (Tisue and Wilensky, 2004; Emonet et al., 2005; Zubler and Douglas, 2009; Koene *et al.*, 2009; Richmond *et al.*, 2010; Collier and North, 2011; Lardon *et al.*, 2011; Rudge *et al.*, 2012; Mirams *et al.*, 2013; Torben-Nielsen and De Schutter, 2014; Kang *et al.*, 2014; Cytowski and Szymanska, 2014; Matyjaszkiewicz *et al.*, 2017; Ghaffarizadeh *et al.*, 2018). In this section, we compare BioDynaMo’s most crucial system properties with prior work.

#### Large-scale model support

The authors of BioCellion (Kang *et al.*, 2014), PhysiCell (Ghaffarizadeh *et al.*, 2018), Timothy (Cytowski and Szymanska, 2014), and Repast HPC (Collier and North, 2011) recognize the necessity for efficient implementations to enable large-scale models. Although these tools can simulate a large number of agents, they do not support neural development. The NeuroMaC neuroscientific simulator (Torben-Nielsen and De Schutter, 2014) claims to be scalable, but the authors do not present performance data and present simulations with only 100 neurons. Therefore, BioDynaMo’s ability to simulate large-scale neural development, which we demonstrate in the results section, is, to our knowledge, unrivaled.

#### General-purpose platform

Many simulators focus on a specific application area: bacterial colonies (Emonet *et al.*, 2005; Matyjaszkiewicz *et al.*, 2017; Rudge *et al.*, 2012; Lardon *et al.*, 2011), cell colonies (Kang *et al.*, 2014; Mirams *et al.*, 2013; Cytowski and Szymanska, 2014), and neural development (Zubler and Douglas, 2009; Koene *et al.*, 2009; Torben-Nielsen and De Schutter, 2014). Pronounced specialization of a simulator may prevent its capacity to adapt to different use cases or simulation scenarios. In contrast, BioDynaMo is a general-purpose platform for agent-based simulations by being both modular and extensible.

#### Quality assurance

Automated software testing is the foundation of a modern development workflow. Unfortunately, several simulation tools (Zubler and Douglas, 2009; Rudge *et al.*, 2012; Lardon *et al.*, 2011; Koene *et al.*, 2009; Torben-Nielsen and De Schutter, 2014; Cytowski and Szymanska, 2014) omit these tests. Mirams *et al.* (2013) recognize this shortcoming and describe a rigorous development workflow in their paper. BioDynaMo has over 280 automated tests which are continuously executed on all supported operating systems to ensure high code quality. BioDynaMo’s open-source code base, tutorials, and documentation not only help users get started, but also enable validation by external examiners.

## 2 Design and implementation

In this section, we present the main simulation concepts of BioDynaMo and describe our approach to achieve modularity, extensibility, and high performance. We provide further information about the biological model, software quality, and features like web-based interactive notebooks, and backups in Supplementary File S1 Section 1.

### 2.1 Simulation concepts

BioDynaMo is implemented in the C++ programming language and supports simulations that follow an agent-based approach. Figure 1 gives an overview of BioDynaMo’s main concepts, while Figure 2 illustrates its object-oriented design.

**Fig. 1.**
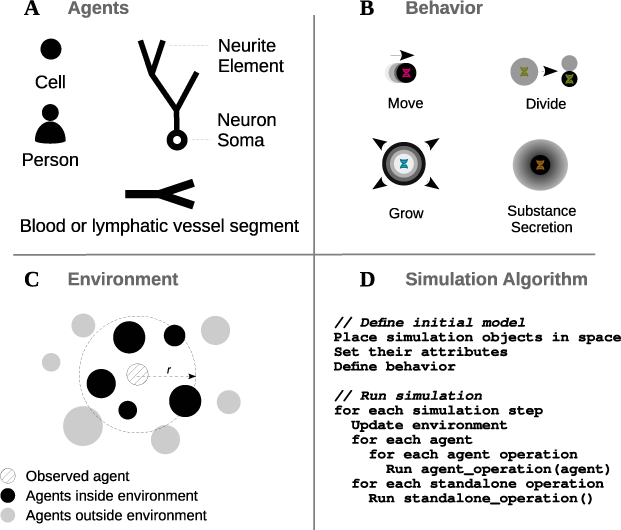
Simulation concepts. Overview of the high-level simulation concepts of BioDynaMo. Agents (A) have their own geometry, behavior (B), and environment (C). (B) Agent behavior is defined in separate classes, which are inserted into agents. A few possible examples for agents and behaviors are displayed. The update of an agent is based on its current state and its surrounding environment. (C) The environment is determined by radius *r* and contains other agents or extracellular substances. The simulation algorithm (D) can be divided into two main parts: the definition of the initial model and execution of the simulation.

**Fig. 2.**
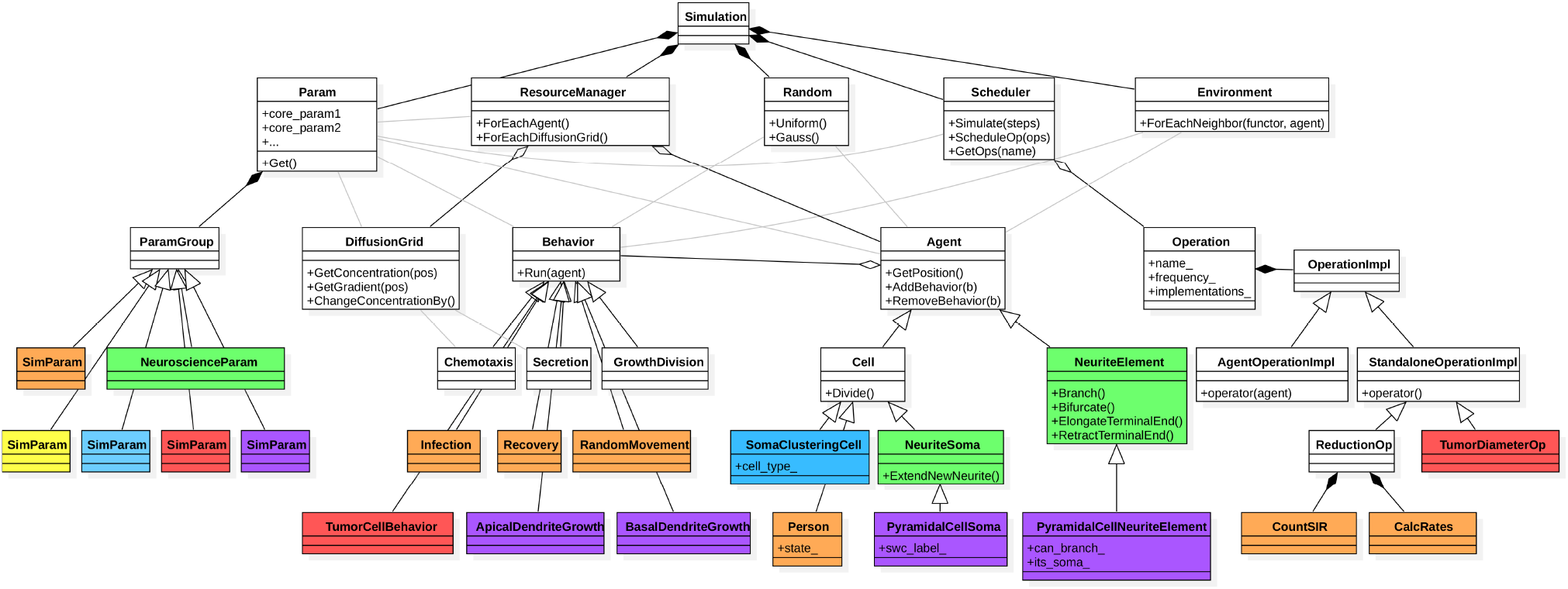
Software design and modularity. Overview of selected classes and functions that are important from the users’s perspective. Classes in white (BioDynaMo core) and green (BioDynaMo’s neuroscience module) are part of the current BioDynaMo installation. The remaining classes illustrate how we extended BioDynaMo to implement the use cases and benchmarks shown in this paper (purple: neuroscience use case, red: oncology use case, orange: epidemiology use case, blue: soma clustering benchmark, yellow: cell proliferation benchmark). A complete list of BioDynaMo classes can be found at https://biodynamo.org/.

A characteristic property of agent-based simulations is the absence of a central organization unit that orchestrates the behavior of all agents. Quite to the contrary, each agent is an autonomous entity that individually determines its behavior. An agent (Figure 1A) has a 3D geometry, behavior, and environment. There is a broad spectrum of entities that can be modeled as an agent. In the results section we show examples where an agent represents a subcellular structure (neuroscience use case), a cell (oncology use case), or an entire person (epidemiology use case). Currently, BioDynaMo supports agents with cylindrical and spherical geometry. Figure 1B shows example agent behaviors such as growth factor secretion, chemotaxis, or cell division. Like genes, behaviors can be activated or inhibited. BioDynaMo achieves this by attaching them to or removing them from the corresponding agent. BioDynaMo simplifies the regulation of behaviors if new agents are created. The user can control if a behavior will be copied to a new agent or removed from the existing agent, based on the event type.

The *Environment* is the vicinity that the agent can interact with (Figure 1C). It comprises other agents and chemical substances in the extracellular matrix. Surrounding agents are, for example, needed to calculate mechanical interactions among agent pairs. BioDynaMo determines the environment based on a uniform grid implementation. The implementation divides the total simulation space into uniform boxes of the same size and assigns agents to boxes based on the center of mass of the agent. Hence, the agents in the environment can be obtained by iterating over the assigned box and all its surrounding boxes (27 boxes in total). The box size is chosen based on the largest agent in the simulation to ensure all mechanical interactions are taken into account.

Currently, the user defines a simulation programmatically in C++ (Figure 1D). There are two main steps involved: initialization and execution. During initialization, the user places agents in space, sets their attributes, and defines their behavior. In the execution step, the simulation engine evaluates the defined model in the simulated physical environment by executing a series of operations. We distinguish between agent operations and standalone operations (Figure 2). At a high level, an agent operation is a function that: (i) alters the state of an agent and potentially also its environment, (ii) creates a new agent, or (iii) removes an agent from the simulation. Examples for agent operations are: execute all behaviors and calculate mechanical forces. The simulation engine executes agent operations for each agent for each time step. Alternatively, standalone operations perform a specific task during one time step and are therefore only invoked once. Examples include the update of substance diffusion and the export of visualization data. Supplementary File S1 Section 1.1.3 contains more details about how operations enable multi-scale simulations.

### 2.2 Modularity

BioDynaMo is a simulation platform that can be used to develop simulations in various computational biology fields (e.g. neuroscience, oncology, epidemiology, etc.). Although agent-based models in these different fields may intrinsically vary, there is a set of functionalities and definitions that they have in common. These commonalities, which consist of simulation and support features, are part of the BioDynaMo core. Simulation features include the physics between cellular bodies, the diffusion of extracellular substances, and basic behavior, such as proliferation and cell death. Support features include visualization, data analysis, plotting, parameter management, simulation backups, etc. Functionalities that are field-specific are separated from the core and are bundled as a separate module. Figure 2 gives an overview of BioDynaMo’s software design. de Montigny *et al.* (2021) demonstrated BioDynaMo’s modularity by coupling it with another simulator to create a hybrid agent-based, continuum-based model.

#### Neuroscience module

The neuroscience module is an example of how to extend functionality in the core to target BioDynaMo to a specific field. The module adds two new agents `NeuronSoma` and `NeuriteElement`, and models behavior like neurite extension from the soma, neurite elongation, and neurite bifurcation. The model closely follows the principles of Cortex3D (Zubler and Douglas, 2009). Neurites are implemented as a binary tree. Each neurite element can have up to two daughter elements. The cylindrical neurite element is approximated as a spring with a point mass on its distal end. These springs are connected to each other to transmit forces along the chain of neurite elements.

#### User-defined components

If the desired functionality is missing, the user can create, extend, or modify agents, behaviors, operations, and other classes as shown in Figure 2. BioDynaMo’s software design focuses on loosely-coupled, well-defined components. This focus not only serves the purpose of creating a clear separation of the functionalities of BioDynaMo, but, perhaps even more significantly, allows users to integrate user-defined components without significant changes to the underlying software architecture. This facilitates collaboration and the creation of an open-model library. We anticipate this library will help researchers in implementing their models more rapidly.

### 2.3 Performance and parallelism

BioDynaMo’s performance is based on the following seven enhancements: (i) The whole simulation engine is parallelized using OpenMP (OpenMP Architecture Review Board, 2015) compiler directives. OpenMP is a good fit since BioDynaMo exploits mostly loop parallelism (see Figure 1A). (ii) To increase the maximum theoretical speedup due to parallel processing (as described by Amdahl’s law (Amdahl, 1967)), we minimize the number of serial code portions in BioDynaMo. (iii) We avoid unnecessary copying of data and optimize data access patterns on machines with non-uniform memory access (NUMA) architecture. Compute nodes with multiple NUMA nodes have different memory access latencies depending on whether a thread accesses local or remote memory. Therefore, we load-balance agents and their environment on available NUMA nodes. We built an optimized iterator over all agents to minimize threads’ memory accesses to non-local memory. This is necessary because OpenMP does not have built-in support for such functionality. (iv) We detect stationary regions within the simulation and skip the expensive collision detection for those agents. (v) We perform just-in-time compilation to give the visualization engine ParaView direct access to Agent attributes. (vi) We develop an optimized memory allocator and concurrent hashmap. (vii) We consider offloading computations to hardware accelerator in our software design (see Figure 2). Our GPU code is implemented in NVidia CUDA and OpenCL and can be executed on graphics cards of different vendors (NVidia, AMD, or Intel). More details on BioDynaMo’s performance enhancements and analyses are beyond this paper’s scope, and we aim to report them in a future publication.

## 3 Results

This section demonstrates BioDynaMo’s capacity to simulate disparate problems in systems biology with simple yet representative use cases in neuroscience, oncology, and epidemiology. Furthermore, we compare BioDynaMo’s performance with an established serial neural simulator (Zubler and Douglas, 2009), analyze its scalability, and quantify the impact of GPU acceleration. For each use case we provide pseudocode for all agent behaviors, a table with model parameters, and more detailed performance results in Supplementary File S1 Section 2.

### 3.1 Neuroscience use case

This example illustrates the use of BioDynaMo to model neurite growth of pyramidal cells using chemical cues. Initially, a pyramidal cell, composed of a 10 *μm* cell body, three 0.5 *μm* long basal dendrites, and one 0.5 *μm* long apical dendrite (all of them considered here as agents), is created in 3D space. Furthermore, two artificial growth factors were initialized, following a Gaussian distribution along the z-axis. The distribution of these growth factors guided dendrite growth and remained unchanged during the simulation.

Dendritic development was dictated by a behavior defining growth direction, speed, and branching behavior for apical and basal dendrites. At each step of the simulation, the dendritic growth direction depended on the local chemical growth factor gradient, the dendrite’s previous direction, and a randomly chosen direction. In addition, the dendrite’s diameter tapered as it grew (shrinkage), until it reached a specified diameter, preventing it from growing any further. The weight of each element on the direction varied between apical and basal dendrites. Apical dendrites were more driven by the chemical gradient and were growing at twice the speed of basal dendrites. On the contrary, basal dendrites were more conservative in their growth direction; the weight of their previous direction was more important. Likewise, branching behavior differed between apical and basal dendrites. In addition to a higher probability of branching (0.038 and 0.006 for apical and basal respectively), apical dendrites had the possibility to branch only on the main branch of the arbor. On the contrary, basal dendrites were only ruled by a simple probability to branch at each time step.

These simple rules gave rise to a straight long apical dendrite with a simple branching pattern and more dispersed basal dendrites, as shown in Figure 3A, similar to what can be observed in real pyramidal cell morphologies as shown in Figure 3B or Spruston (2008) (Figure 1A CA1). Using our growth model, we were able to generate a large number of various realistic pyramidal cell morphologies. We used a publicly available database of real pyramidal cells coming from (Mellström *et al.*, 2016) for comparison and parameter tuning. Two measures were used to compare our simulated neurons and the 69 neurons composing the real morphologies database: the average number of branching points, and the average length of dendritic trees. No significant differences were observed between our simulated neurons and the real ones (*p* < 0.001 using a T-test for two independent samples). These results are shown in Figure 3D. The simulation of the pyramidal cell growth consisted of 361 lines of C++ code.

**Fig. 3.**
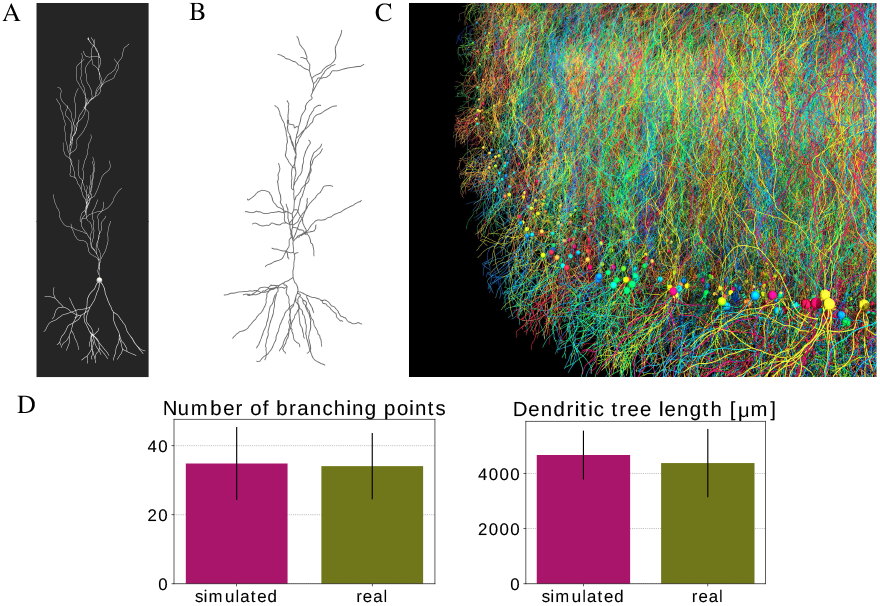
Pyramidal cell simulation. (A) Example pyramidal cell simulated with BioDynaMo. (B) Real neuron (R67nr67b-CEL1.CNG) taken from (Mellström et al., 2016) and visualized with https://neuroinformatics.nl/HBP/morphology-viewer/ (C) Large-scale simulation. The model started with 5000 initial pyramidal cell bodies and contained 9 million agents after simulating 500 iterations. Simulation execution time was 46 seconds on a server with 72 CPU cores. (D) Morphology comparison between simulated neurons and experimental data from (Mellström et al., 2016). Error bars represent the standard deviation. (A,C) A video is available in Supplementary Information.

Figure 3C shows a large scale simulation incorporating 5000 neurons similar to the one described above, and demonstrates the potential of BioDynaMo for developmental, anatomical, and connectivity studies in the brain. This simulation contained 9 million agents. These 500 iterations correspond to approximately three weeks of pyramidal cell growth in the rat.

### 3.2 Oncology use case

In this section, we present a tumor spheroid simulation to replicate in vitro experiments from (Gong *et al.*, 2015). Tumor spheroid experiments are typically employed to investigate the pathophysiology of cancer, and are also being used for pre-clinical drug screening (Nunes *et al.*, 2019). Here we considered three in vitro test cases using a breast adenocarcinoma MCF-7 cell line (Gong *et al.*, 2015) with different initial cell populations (2000, 4000, and 8000 MCF-7 cells). Our goal was to simulate the growth of this mono cell culture embedded in a collagenous (extracellular) matrix. This approach, as opposed to a free suspension one, incorporates cell-matrix interactions to mimic the tumor-host environment.

Initially, cancer cells (agents) were clustered in a spherical shape around the origin with a diameter of 310, 380, or 460 micrometers. The three-dimensional extracellular matrix (ECM) was represented in our simulations as a 8 mm^3^ cube. The fundamental cellular mechanisms modeled here include cell growth, cell duplication, cell migration, and cell apoptosis. A single behavior governed all these processes. The cell growth rate was derived from the published data (Sutherland *et al.*, 1983), while cell migration (cell movement speed), cell survival, and apoptosis were fine-tuned after trial and error testing. Since the in vitro study considered the same agarose gel matrix composition among the experiments, the BioDynaMo model assumes identical parameters for the cell–matrix interactions in the simulations. Considering the homogeneous ECM properties, tumor cell migration was modeled as Brownian motion.

The in vitro experiments showed that instantaneous spheroid growth was hindered by the compression of the surrounding agarose gel matrix (see Figure 4A), owing to cell reorganization at the onset of the cancer mass implantation into the gel. As a result, the tumor spheroid diameter was initially decreasing. However, the present simulation example focuses modeling the growth of the spheroid after it had set in the agarose gel matrix. Therefore, as shown in Figure 4A, BioDynaMo simulations are set to start on day two or three.

**Fig. 4.**
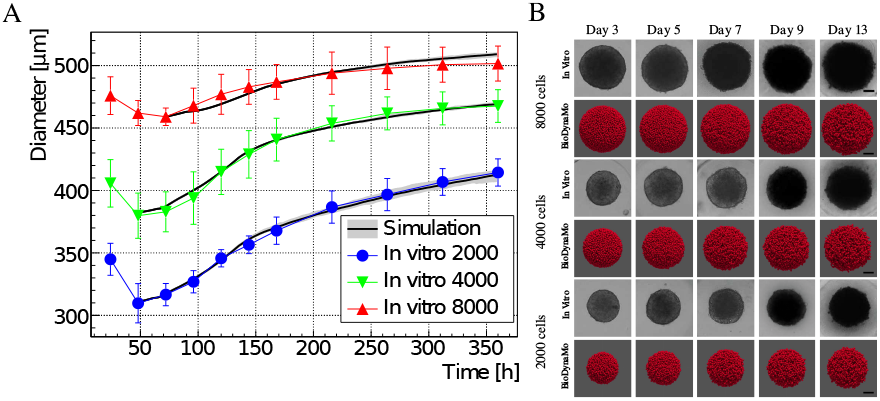
Comparison between in vitro MCF-7 tumor spheroid experiments and our in silico simulations using BioDynaMo. (A) Human breast adenocarcinoma tumor spheroid (MCF-7 cell line) development during a 15 day period, where different initial cell populations were considered (see Fig 3 in (Gong et al., 2015)). Error bars denote standard deviation to the experimental data. The mean of the in silico results is shown as a solid black line with a grey band depicting minimum and maximum observed value. (B) Qualitative comparison between the microscopy images and simulation snap-shots is shown in the three boxes. Scale bars correspond to 100μm. A video is available in Supplementary Information.

The in vitro experiments from (Gong *et al.*, 2015) and the simulations using BioDynaMo are depicted in Figure 4. Each line plot in Figure 4A compares the mean diameter between the experiments and the simulations over time, which demonstrates the validity and accuracy of BioDynaMo. The diameter of the spheroids in the simulations were deducted from the volume of the convex hull that enclosed all cancer cells. The in vitro experiments used microscopy imaging to measure the spheroid’s diameters (Gong *et al.*, 2015). Figure 4B compares snapshots of the simulated tumor spheroids (bottom row) against microscopy images of in vitro spheroids (top row) at different time points. The spheroid’s morphologies between the in vitro experiments and the BioDynaMo simulations are in excellent agreement.

The example has 424 lines of C++ code, including the generation of the plot shown in Figure 4A. Running one simulation took 0.98–3.39s on a laptop and 1.24–4.16s on a server, both using one CPU core.

### 3.3 Epidemiology use case

This section presents an agent-based model that describes the spreading of infectious diseases between humans. The model divides the population into three groups: susceptible, infected, and recovered (SIR) agents. We compare our simulation results with the solution of the original SIR model from Kermack *et al.* (1927), which used the following three differential equation to describe the model dynamics: *dS/dt* = *-βSI/N*, *dI/dt* = *βSI/N* – *γI*, and *dR/dt* = *γI. S, I*, and *R* are the number of susceptible, infected, and recovered individuals, *N* is the total number of individuals, *β* is the mean transmission rate, and γ the recovery rate.

For our agent-based implementation (Figure 5C) we created a new agent (representing a person) that encompasses three new behaviors, and extended an operation to count the number of agents in each group (see Figure 2). Agents were randomly distributed in space and have three behaviors. Infection. A susceptible agent became infected with the infection probability if an infected agent was within the infection radius. Recovery. An infected agent recovered with the recovery probability at every time step. Random movement. All agents moved randomly in space. The absolute distance an agent may travel in every time step is limited.

**Fig. 5.**
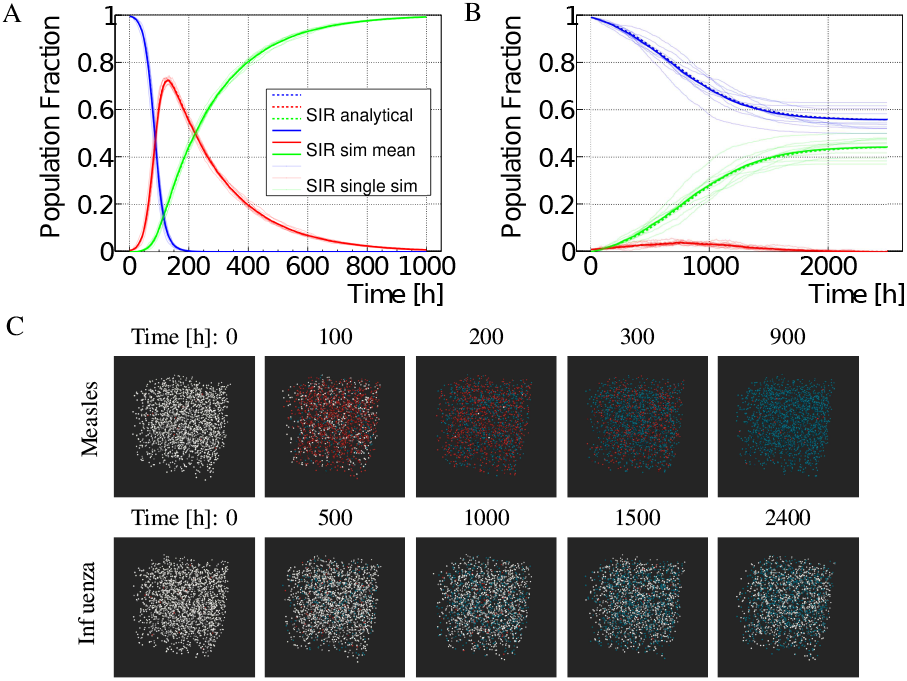
Measles and seasonal influenza SIR model results. (A,B) Comparison between agent-based (solid lines) and analytical (dashed lines) model for measles (A) and seasonal influenza (B). The agent-based simulation was repeated ten times. The individual simulation results are shown as thin solid lines. The bold solid line represents the mean from all simulations. The legend is shared between the two plots. (C) Visualization of the measles and influenza model for different time steps in 3D space. Susceptible persons are shown in white, infected persons in red, and recovered persons in blue. Persons move randomly and follow the rules for infection and recovery.

In this agent-based model, the speed at which an infectious disease spreads depended on: the infection probability, the number of contacts each agent has with other agents, and the recovery rate. The number of contacts in turn depended on the infection radius, the maximum distance an agent may travel, and the density of agents in the simulation space.

We selected two infectious diseases with different characteristics to verify our model: measles and seasonal influenza. We obtained values for the basic reproduction number *R*_0_ and recovery duration *T_R_* from the literature (Measles: *R*_0_ = 12.9, *T_R_* = 8 days (Guerra *et al.*, 2017; World Health Organization, 2020), Influenza: *R*_0_ = 1.3, *T_R_* = 4.1 days (Chowell *et al.*, 2008)) and determined the parameters *β* and *γ* for the analytical model, based on *R*_0_ = *β/γ* and *γ* = 1/*T_R_*. For the agent-based model we set the recovery probability to γ, and placed 2000 susceptible agents and a few infected agents randomly in a cubic space with length 100. The remaining parameters (infection radius, infection probability, and maximum movement in one time step) were determined using particle swarm optimization (Kennedy and Eberhart, 1995). Figure 5 shows that the agent-based model is in excellent agreement with the equation-based approach from (Kermack *et al.*, 1927) for measles and influenza.

The example has 566 lines of C++ code, including the generation of the plot shown in Figure 5. Running one simulation took 0.59–1.59s using one CPU core.

### 3.4 Performance

Efficient usage of computing resources is paramount for large-scale simulations with billions of agents, reduced computational costs, and low energy footprint. To this end, we quantify the performance of BioDynaMo with three simulations: cell growth and division, soma clustering, and pyramidal cell growth. These simulations have different properties and are, therefore, well suited to evaluate BioDynaMo’s simulation engine under a broad set of conditions. Supplementary File S1 Section 2.2 contains more details about these benchmarks.

First, to demonstrate the performance improvements against established agent-based simulators, we compared BioDynaMo with Cortex3D (Zubler and Douglas, 2009). Cortex3D has the highest similarity in terms of the underlying biological model out of all the related works presented in Section 1.1. More specifically, BioDynaMo and Cortex3D use the same method to determine mechanical forces between agents and the same model to grow neural morphologies. This makes Cortex3D the best candidate with which to compare BioDynaMo and ensure a fair comparison. Figure 6A shows the speedup of BioDynaMo for the three simulations. We observed a significant speedup between 18 and 78 ×. Note that we set the number of threads available to BioDynaMo to one since Cortex3D is not parallelized. The speedup was larger, when the simulation was more dynamic or more complex.

**Fig. 6.**
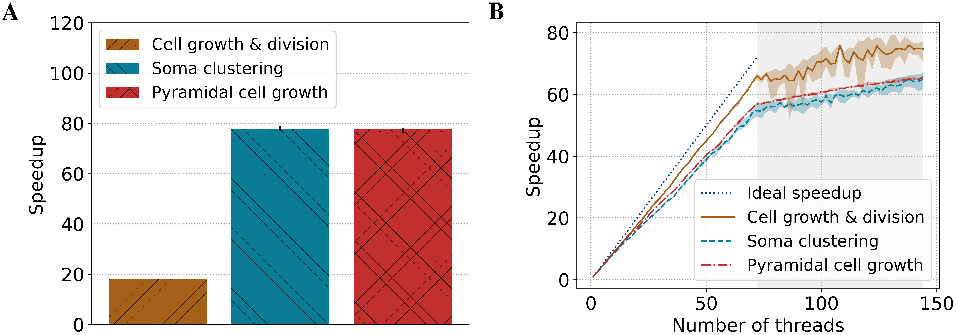
BioDynaMo performance analysis. (A) Speedup of BioDynaMo compared to Cortex3D. (B) Strong scaling behavior of BioDynaMo on a server with 72 physical cores, two threads per core, and four NUMA domains. The grey area highlights hyper-threads.

Second, to evaluate the scalability of BioDynaMo, we measured the simulation time with an increasing number of threads. We increased the number of agents used in the comparison with Cortex3D and reduced the number of simulation timesteps to 10. Figure 6B shows the strong scaling analysis. All simulation parameters were kept constant, and the number of threads was increased from one to the number of logical cores provided by the benchmark server. The maximum speedup ranged between 65× and 75×, which corresponds to a parallel efficiency of 0.90 and 1.04. Performance improved even after all physical cores were utilized and hyper-threads were used. Hyper-threads are highlighted in gray in Figure 6B. We want to emphasize that even the pyramidal cell growth benchmark scaled well, despite the challenges of synchronization and load imbalance.

Third, we evaluated the impact of calculating the mechanical forces on the GPU using the cell growth and division, and soma clustering simulations. We excluded the pyramidal cell growth simulation because the current GPU kernel does not support cylinder geometry yet. The benchmarks were executed on System C (see Supplementary File S1 Table 4), comparing an NVidia Tesla V100 GPU with 32 CPU cores (64 threads). We observed a speedup of 1.27 × for cell growth and division, and 5.04 × for soma clustering. The speedup correlated with the number of collisions in the simulation. The computational intensity is directly linked with the number of collisions between agents.

In summary, in the scalability test, we observed a minimum speedup of 65 ×. Furthermore, we measured a minimum speedup of 18× comparing BioDynaMo with Cortex3D both using a single thread. Based on these two observations, we conclude that on System A (see Supplementary File S1 Table 4) BioDynaMo is more than three orders of magnitude faster than Cortex3D.

Based on these speedups, we executed the neuroscience, oncology, and epidemiology use cases with one billion agents. Using all 72 physical CPUs on System B (see Supplementary File S1 Table 4), we measured a runtime of 1 hour 37 minutes, 6 hours 49 minutes, and 3 hours 54 minutes, respectively. One billion agents, however, are not the limit. The maximum depends on the available memory and accepted execution duration. To be consistent across all use cases and keep our pipeline’s total execution time better manageable, we decided to run these benchmarks with one billion agents. Table 5 in Supplementary File S1 shows that available memory would permit an epidemiological simulation with three billion agents. With enough memory, BioDynaMo is capable of supporting hundreds of billions of agents.

## 4 Discussion

This paper presented BioDynaMo, a novel open-source platform for agent-based simulations. Its modular software architecture allows researchers to implement models of distinctly different fields, of which neuroscience, oncology, and epidemiology were demonstrated in this paper. Although the implemented models follow a simplistic set of rules, the results that emerge from the simulations are prominent and highlight BioDynaMo’s capabilities. We do not claim that these models are novel, but we rather want to emphasize that BioDynaMo enables scientists to (i) develop models in various computational biology fields in a modular fashion, (ii) obtain results rapidly with the parallelized execution engine, (iii) scale up the model to billions of agents on a single server, and (iv) produce results that are in agreement with validated experimental data. Although BioDynaMo is modular, we currently offer a limited number of ready-to-use simulation primitives. We are currently expanding our library of agents and behaviors to facilitate model development beyond the current capacity.

Ongoing work uses BioDynaMo to gain insights into retinal development, cryopreservation, multiscale (organ-to-cell) cancer modelling, COVID-19 spreading in closed environments, radiation-induced tissue damage, and more. Further efforts focus on accelerating drug development by replacing in vitro experiments with in silico simulations using BioDynaMo.

Our performance analysis showed improvements of up to three orders of magnitude over state-of-the-art baseline simulation software, allowing us to scale up simulations to an unprecedented number of agents. To the best of our knowledge, BioDynaMo is the first scalable simulator of neural development with cellular interactions that scales to more than one billion agents. The same principles used to model axons and dendrites in the neuroscience use case could also be applied to simulate blood and lymphatic vessels.

We envision BioDynaMo to become a valuable tool in computational biology, fostering faster and easier simulation of complex and large-scale systems, interdisciplinary collaboration, and scientific reproducibility.

## Acknowledgments

We want to thank Giovanni De Toni for his work on the BioDynaMo build system.

